# Knowledge, attitudes and awareness of the human papillomavirus among health professionals in New Zealand

**DOI:** 10.1101/317248

**Authors:** Susan M. Sherman, Karen Bartholomew, Hayley J. Denison, Hersha Patel, Esther L. Moss, Jeroen Douwes, Collette Bromhead

## Abstract

**Background:** Human papillomavirus (HPV) is a common sexually transmitted infection that is implicated in 99.7% of cervical cancers and several other cancers that affect both men and women. Despite the role that HPV plays in an estimated 5% of all cancers and the evolving role of HPV vaccination and testing in protecting the public against these cancers, preliminary research in New Zealand health care professionals suggest knowledge about HPV may not be sufficient.

**Methods:** A total of 230 practice nurses, smear takers and other clinical and laboratory staff who attended a range of training events completed a cross-sectional survey between April 2016 and July 2017. The survey explored four broad areas: demographics and level of experience, HPV knowledge (general HPV knowledge, HPV triage and test of cure (TOC) knowledge and HPV vaccine knowledge), attitudes towards the HPV vaccine and self-perceived adequacy of HPV knowledge.

**Results:** The mean score on the general HPV knowledge questions was 13.2 out of 15, with only 25.2% of respondents scoring 100%. In response to an additional question, 14.0% thought (or were unsure) that HPV causes HIV/AIDS. The mean score on the HPV Triage and TOC knowledge questions was 7.4 out of 10, with only 9.1% scoring 100%. The mean score on the HPV vaccine knowledge questions was 6.0 out of 7 and 44.3% scored 100%. Only 62.6% of respondents agreed or strongly agreed that they were adequately informed about HPV, although 71.8% agreed or strongly agreed that they could confidently answer HPV-related questions asked by patients. Multivariate analyses revealed that knowledge in each domain predicted confidence in responding to patient questions. Furthermore, the number of years since training predicted both HPV knowledge and Triage and TOC knowledge.

**Discussion:** Although overall level of knowledge was adequate, there were significant gaps in knowledge, particularly about the role of HPV testing in the New Zealand National Cervical Screening Programme. More education is required to ensure that misinformation and stigma do not inadvertently result from interactions between health practitioners and the public.

## Introduction

Human papillomavirus (HPV) is responsible for 99.7% of cases of cervical cancer along with some head and neck, penile and anal cancers. There are approximately 150 new diagnoses and 50 deaths from cervical cancer in New Zealand (NZ) every year^1^, while head and neck cancers attributable to HPV are increasing in both men and women with 94 new cases and 43 deaths estimated for 2012^2^. In addition, there are longstanding ethnic inequalities in cervical cancer incidence and mortality, and cervical screening coverage remains low (and cancer incidence and mortality high) for indigenous Māori women as well as Pacific women^3^.

The NZ National Cervical Screening Programme (NCSP), which was established in 1990^4^, recommends 3-yearly routine screening with liquid-based cytology (LBC) for 20–69 year-old women, with HPV triage testing for low grade (ASC-US/LSIL) cytology in women 30+ years. The programme also recommends testing of cure following treatment for a high-grade lesion^4^, with a modified version of the Bethesda System used for cytology classification. However, from late 2018 the NCSP will introduce HPV testing as the primary screening test for women aged 25-68 years on a 5 yearly basis^5^.

To reduce infection with high-risk types of HPV and its related cancers, the NZ National HPV Immunisation Programme was introduced in September 2008, offering free HPV vaccination (Gardasil^®^, Merck) for females born in 1990 or later. School-based immunisation for 12–13 year-old girls commenced in most regions in 2009^1^ and the three-dose coverage achieved by the program in cohorts born in 1991–2002 reached approximately 48–66% nationwide^1^. In January 2017, the free programme was extended to boys and young men, the upper age for free vaccination was increased to 26 years, a two-dose schedule was implemented for individuals aged 14 and under, and the vaccine used was changed to nonavalent Gardasil 9 (Merck)^1^.

Previous research has shown that the public is not well informed about HPV. For example, in a survey of 200 NZ university health science students (mean age 19.8 years), 50.8% were both unaware of the sexual transmission of HPV and unwilling to accept a free HPV vaccine, highlighting the need for education in this age group^6^.

Furthermore, it has been shown that there is considerable stigma associated with a diagnosis of HPV^7, 8^. For example, in a qualitative study, McCaffrey et al., found that HPV positive women^7^ reported levels of stigma and anxiety suggesting that “testing positive for HPV was associated with adverse social and psychological consequences that were beyond those experienced by an abnormal smear alone “ (p173). Daley et al., found that younger age and less education were associated with more negative emotions (e.g., anger, shock and worry) and stigma beliefs (e.g., feeling ashamed, guilty and unclean) in HPV positive women^8^. In addition, a survey of Hong Kong Chinese health care providers exploring levels of knowledge about HPV and attitudes revealed that more knowledge about HPV predicted less stigmatising attitudes from health care providers^9^. Since NZ women will, from late 2018, be tested for HPV every 5 years, it is crucial that the health care professionals who administer the screening programme, and who will be best placed to counsel women who receive a positive HPV test, are themselves well informed about HPV. This will also likely reduce inequalities in screening uptake. In particular, previous research concerning the diagnosis of depression identified that effective patient-clinician communication and an established relationship in which the patient trusted the general practitioner (GP) were especially important for Māori patients^10^.

Previous research exploring the knowledge of GPs and practice nurses (PNs) in Christchurch, New Zealand about HPV used 5 questions as part of a larger survey exploring attitudes towards HPV vaccination^11^. Whilst performance across the 5 questions was reasonable, there was uncertainty as indicated by the number of ‘not sure’ responses, as well as some variability across questions. For example, while more than 90% of GPs and PNs knew that HPV vaccination would not eliminate the need for cervical screening, only 33% of GPs and 7% of PNs knew that anogenital warts caused by HPV 6 and 11 are not a precursor to cervical cancer. Only half of GPs and 42% of PNs knew that most HPV infections will clear without medical treatment and a quarter of GPs and nearly a third of PNs did not know, or were unsure, whether persistent HPV was a necessary cause of cervical cancer.

A recent study in the UK conducted a more detailed online survey of PNs in Leicestershire^12^. While overall awareness of the basic facts about HPV was adequate, there were some worrying gaps in knowledge, for example nearly 10% of PNs did not know that HPV causes cervical cancer and 63% believed that HPV requires treatment. Some of the PNs reported not feeling adequately informed about HPV and that training provision could be improved. There was no correlation found between self-perceived adequacy of knowledge and HPV knowledge scores.

To our knowledge there are no studies exploring what primary care staff such as GPs, PNs and smear takers in NZ know about HPV since 2009. In light of the recent changes to the immunisation programme and the forthcoming changes to the NSCP, it is important to benchmark what nurses and smear takers understand about HPV, whether they feel well informed and assess any training needs they might identify.

## Methods

An anonymous cross-sectional survey was conducted between April 2016 and July 2017. GPs, practice nurses, smear takers and other clinical and laboratory staff who attended a variety of training events in Auckland District Health Board (DHB) and Waitemata DHB catchment areas were invited to complete the paper-based survey.

The survey was taken from Patel et al.,^12^ who had incorporated most of the items from Waller et al.,^13^ and was adapted by adding back in a question about HPV and HIV/AIDS from Waller et al., and by changing some wording to make the terminology or protocols New Zealand-specific. The overall face validity of the adapted instrument was confirmed by peer review by members of the immunisation team in Auckland and cervical screening expert doctors/nurse practitioners.

The final survey explored four broad categories: demographics and level of experience; HPV knowledge (general HPV knowledge, HPV triage and test of cure (TOC) knowledge and HPV vaccine knowledge), which were assessed using a true, false, don’t know format; and attitudes towards the HPV vaccine and self-perceived adequacy of HPV knowledge, which were assessed using 5-point Likert scales (the survey is publicly available here: osf.io/ub7g2, DOI 10.17605/OSF.IO/UB7G2).

### Statistical analyses

Demographic factors included age, profession and years since HPV training. For analyses, profession was collapsed into four categories (nurse; general practitioner (GP); colposcopy, which included colposcopists and colposcopy nurses; and laboratory staff and other), and years since HPV training was collapsed into 3 categories (never; ≤ 1 year; > 1 year).

Factors affecting HPV knowledge were explored using ordinal regression analysis. First, univariate regression analyses were conducted to identify variables associated with knowledge. Then, variables with a p value <0.05 were entered into a multivariate model to estimate the association between these predictors and knowledge score, while controlling for potential confounders.

Factors affecting self-perceived adequacy of HPV knowledge were also explored, using binary logistic regression. Feeling adequately informed and feeling confident in answering patient questions were converted from 5-point Likert scales to binary variables of yes (strongly agree, agree) and no or undecided (strongly disagree, disagree or undecided) as the dependent variables. Again, predictor variables which were associated with the dependent variables in univariate analyses were entered into a multivariate model to estimate the association between knowledge score and self-perceived adequacy while controlling for potential confounders.

## Results

A total of 234 individuals completed the survey. The data for four individuals were removed, as there were large sections that had been left unanswered. The majority (212) were female and one person did not identify a gender. Of the 230 participants, 107 had never taken a smear. For the 123 who had, the years of experience ranged from 0.1 to 42 years (mean 8.7 years, median 6.5 years). Details about age categories, profession and date of most recent training, if any, are presented in Table 1.

**Table 1.**
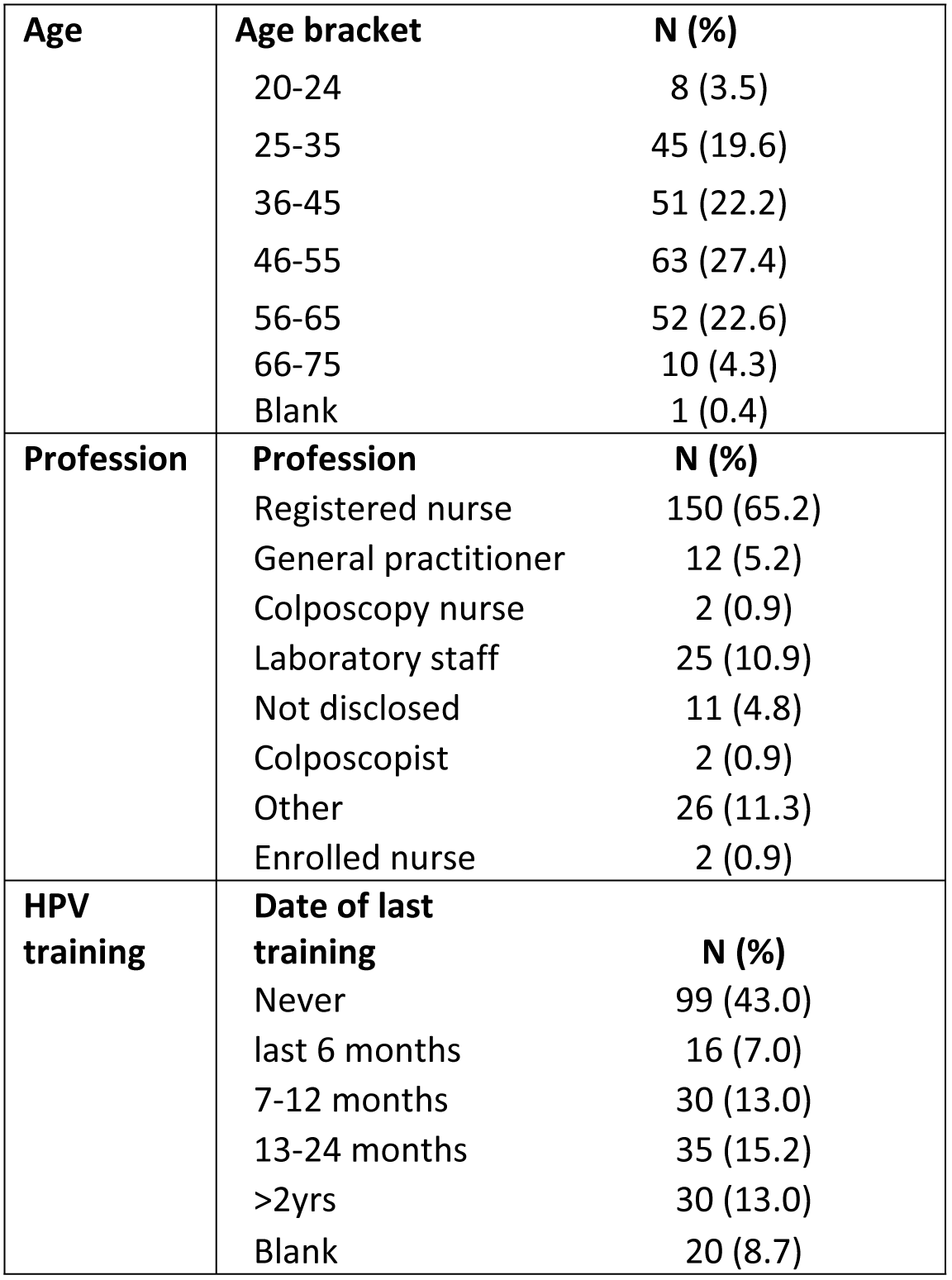
Participant characteristics.

### General HPV knowledge

Out of a maximum knowledge score of 15 (see individual questions in Table 2 and excluding the question about HIV/AIDS), the mean score achieved by participants was 13.2 (standard deviation (SD) 2.0) and the median score was 14 (range 0-15, interquartile range (IQR) 13-15), with 25.2% (N=58) achieving 100%. One individual did not answer any questions correctly.

**Table 2.**
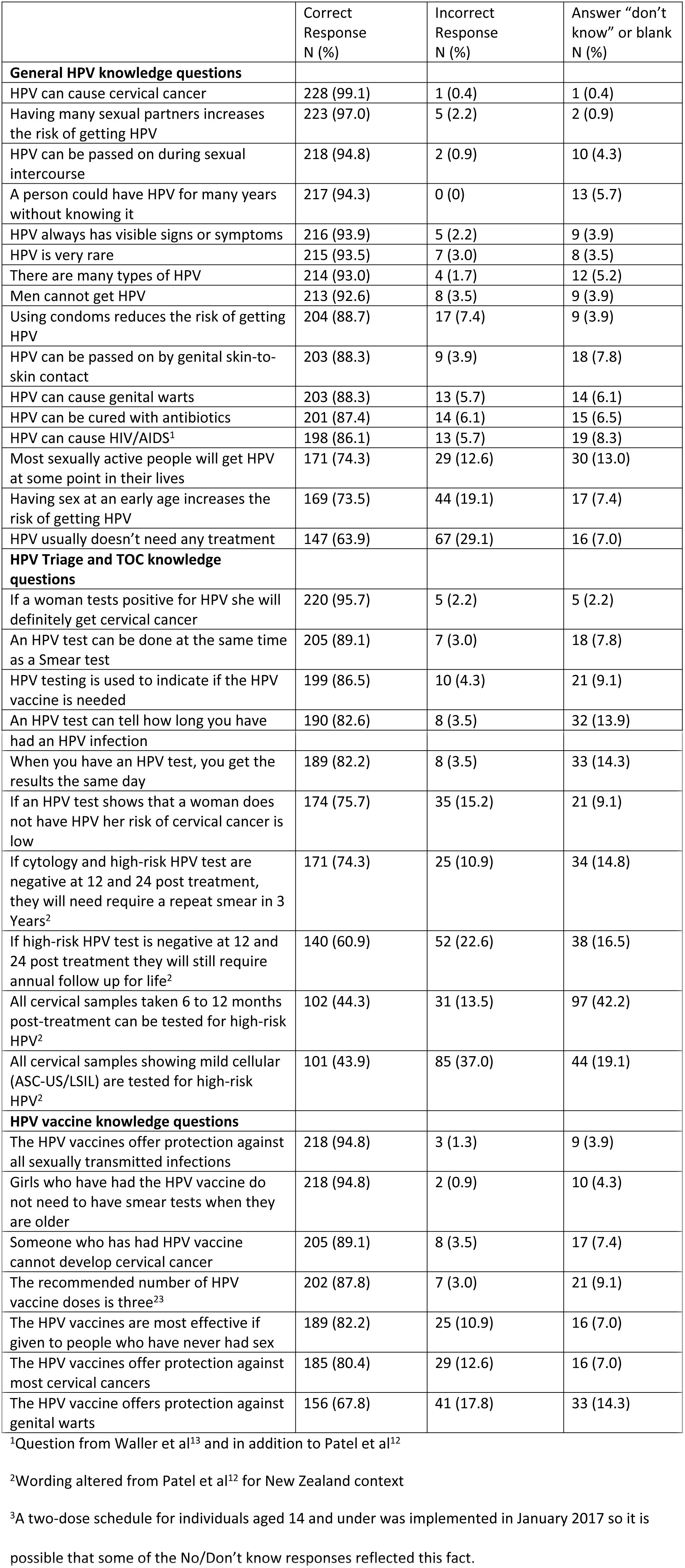
HPV and vaccine knowledge questions

The following questions were most often answered incorrectly: HPV usually doesn’t need any treatment (36.1% answered incorrectly or weren’t sure); Having sex at an early age increases the risk of getting HPV (26.5%); Most sexually active people will get HPV at some point in their lives (25.7%). In addition, more than 10% of individuals incorrectly thought (or were not sure) that HPV cannot be passed on by genital skin-to-skin contact, that HPV does not cause genital warts, that using condoms does not reduce the risk of getting HPV and that HPV can be cured with antibiotics.

Following Waller et al.,^13^ the item about HIV/AIDS was analysed separately from the rest of the questions. In total, 86.1% of respondents correctly identified that HPV does not cause HIV/AIDS.

### HPV Triage and TOC knowledge

Out of a maximum knowledge score of 10 (see individual questions in Table 2), the mean score achieved by the participants was 7.4 (SD 2.1) and the median score was 8 (range 0-10, IQR 6-9), with 9.1% (N=21) achieving 100%. Three individuals had no correct answers.

The following questions were answered incorrectly most often: All cervical samples showing mild cellular (ASC-US/LSIL) are tested for high-risk HPV (56.1% answered incorrectly or were not sure); All cervical samples taken 6 to 12 months post-treatment can be tested for high-risk HPV (55.7%); If high-risk HPV test is negative at 12 and 24 post treatment they will still require annual follow up for life (39.1%); If cytology and high-risk HPV test are negative at 12 and 24 post treatment, they will require a repeat smear in 3 Years (25.7%). In addition, more than 10% of individuals incorrectly thought (or weren’t sure) that an HPV test can tell how long a person has had an HPV infection; an HPV test cannot be done at the same time as a Smear test; HPV testing is used to indicate if the HPV vaccine is needed; when an HPV test has been done that the results are available the same day; If an HPV test shows that a women does not have HPV her risk of cervical cancer is not low.

### HPV vaccine knowledge

Out of a maximum knowledge score of 7 (see individual questions in Table 2), the mean score achieved by the participants was 6.0 (SD 1.2) and the median score was 6 (range 0-7, IQR 5-7), with 44.3% (N=102) achieving 100%. One individual had no answers correct.

The following questions were answered incorrectly most often: The HPV vaccine offers protection against genital warts (32.2% answered incorrectly or weren’t sure); The HPV vaccines offer protection against most cervical cancers (19.6%); The HPV vaccines are most effective if given to people who have never had sex (17.8%). In addition, more than 10% of participants incorrectly thought (or weren’t sure) that the recommended number of HPV vaccine doses was not three and that someone who has had the HPV vaccine cannot develop cervical cancer.

### Factors influencing level of HPV knowledge

Table 3 shows the effect of predictors on the three types of knowledge, both unadjusted (‘crude’) and adjusted for the other covariates (‘full model’). Having ever taken a smear was significantly positively associated with all three types of knowledge when entered into the model as the only predictor. However, when adjusting for the other predictors, the association with having ever taken a smear was attenuated for all knowledge types and only remained significantly associated with Triage and TOC knowledge score (where those who had ever taken a smear were 4 times (OR 4.1, 95% CI 2.1 – 8.1) more likely to have a higher knowledge score than those who had not taken a smear, p < 0.01).

**Table 3.**
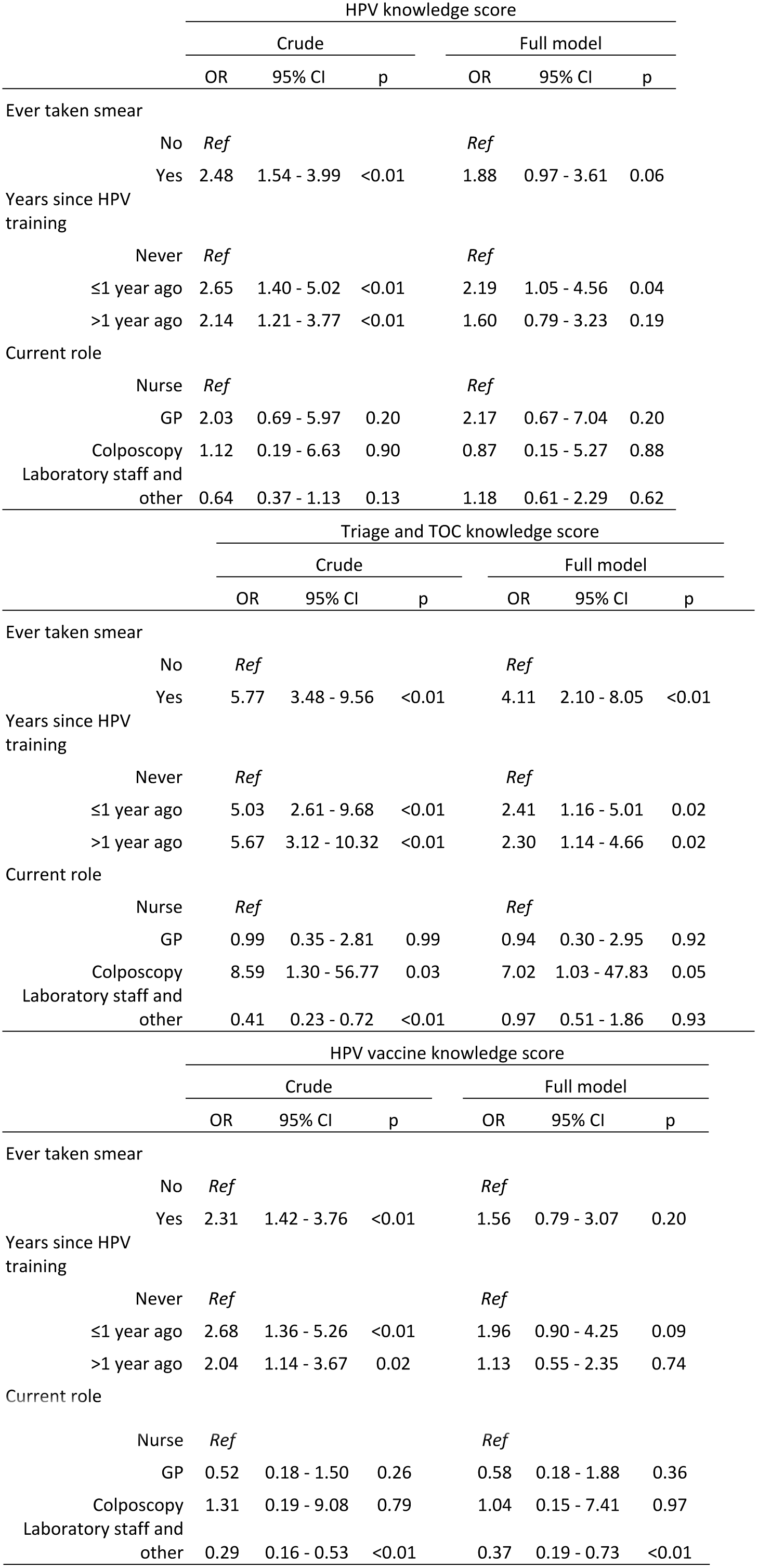
Ordinal regression of predictors of knowledge

Years since HPV training was also associated with knowledge level in univariate analysis, where those who had had training (either ≤ 1 year ago or > 1 year ago) were more likely to have a higher knowledge score than those who had never had HPV training, across all types of knowledge. The association was more pronounced for those who had had more recent training (≤ 1 year ago) than for those who had training longer ago (> 1 year ago), as expected. The association was attenuated when taking into other predictors on knowledge. However, having had HPV training ≤ 1 year ago compared to never remained significantly independently predictive of HPV knowledge score and Triage and TOC knowledge score; having had training > 1 year ago compared to never also remained significantly independently predictive of Triage and TOC knowledge score. Years since training was not predictive of HPV vaccine knowledge score after adjustment for the other predictors.

Current role was not associated with HPV knowledge score in univariate or multivariate analyses. However, current role was associated with the Triage and TOC knowledge score, where, even after adjustment for the other predictors, those who worked in colposcopy were 7 times (95% CI 1.0 – 47.8) more likely to have a higher knowledge score than nurses (who were the reference category) (p = 0.05). However, the number of colposcopy workers was very small (n=4), so this result should be interpreted with caution. Those that were classed as laboratory staff or other were less likely to have higher Triage and TOC knowledge scores in the univariate analyses, but this association disappeared after adjustment for the other predictors. The laboratory staff and other group were also more likely to have lower HPV vaccine knowledge scores than nurses in both univariate (OR 0.3, 95% CI 0.2 – 0.5, p < 0.01) and multivariate (OR 0.4, 95% CI 0.2 – 0.7, p < 0.01) models.

The effect of age on knowledge score was explored in univariate analysis as a potential predictor, but was not associated with scores for any of the three knowledge types (data not shown), so was not included in the multivariate analysis.

### Attitudes towards HPV vaccine

Of all respondents, 95.6% (N=220) agreed or strongly agreed that they would recommend the HPV vaccine (Table 4), with a further 3.5% (N=8) undecided (there were 2 (0.9%) blank responses).

**Table 4.** Attitudes and adequacy questions

|  | Response N (%) |  |  |  |  |  |
| --- | --- | --- | --- | --- | --- | --- |
|  | Strongly agree | Agree | Undecided | Disagree | Strongly disagree | Blank |
| Would you recommend the HPV vaccine? | 179 (77.8) | 41 (17.8) | 8 (3.5) | 0 | 0 | 2 (0.9) |
| Do you think that the vaccine should be offered to men/boys as well? | 168 (73.0) | 47 (20.4) | 12 (5.2) | 1 (0.4) | 0 | 2 (0.9) |
| Do you feel adequately informed about HPV? | 30 (13.0) | 114 (49.6) | 52 (22.6) | 27 (11.7) | 3 (1.3) | 4 (1.7) |
| Are you able to confidently answer HPV related questions asked by patients? | 37 (16.1) | 128 (55.7) | 41 (17.8) | 15 (6.5) | 4 (1.7) | 5 (2.2) |

In total, 93.4% (N=215) respondents agreed or strongly agreed that men/boys should be offered the vaccine (Table 4), with 5.2% (N=12) undecided and 0.4% (N=1) in disagreement (there were 2 (0.9%) blank responses).

### Self-perceived adequacy of HPV knowledge

Only 62.6% (N=144) respondents agreed or strongly agreed that they were adequately informed about HPV (see Table 4), 22.6% (N=52) were undecided, while 13.0% (N=30) disagreed or strongly disagreed (there were 4 (1.7%) blank responses).

Despite this, 71.8% (N=165) respondents agreed or strongly agreed that they could confidently answer HPV related questions asked by patients (see Table 4). A further 17.8% (N=41) were undecided and 8.2% (N=19) disagreed or strongly disagreed (there were 5 (2.2%) blank responses).

Independent t-tests confirmed that the knowledge scores for general HPV knowledge, triage and test of cure knowledge and HPV vaccine knowledge were all significantly higher for those participants who felt they were adequately informed than in those who did not feel they were or who were unsure (p<0.01). The same was found for the question about feeling confident in answering patient questions.

Feeling adequately informed and feeling confident in answering patient questions were both related to having ever taken a smear, years since training, and to a much lesser extent, current role (data not shown). Therefore, the relationship between self-perceived adequacy and knowledge was explored further in multivariate analysis using binary logistic regression (Table 5).

**Table 5.** Logistic regression of the effect of knowledge on feeling adequately informed / confident in answering patient questions

|  | Feeling adequately informed |  |  |  |  |  |
| --- | --- | --- | --- | --- | --- | --- |
|  | Crude |  |  | Adjusted for ever taken a smear, years since training and current role |  |  |
|  | OR | 95% CI | p | OR | 95% CI | p |
| HPV knowledge | 1.33 | 1.12 – 1.58 | <0.01 | 1.15 | 0.95 – 1.39 | 0.15 |
| Triage and TOC knowledge | 1.31 | 1.13 – 1.51 | <0.01 | 1.05 | 0.88 – 1.25 | 0.61 |
| HPV vaccine knowledge | 1.59 | 1.25 – 2.03 | <0.01 | 1.53 | 1.14 – 2.04 | <0.01 |

|  | Feeling confident in answering patient questions |  |  |  |  |  |
| --- | --- | --- | --- | --- | --- | --- |
|  | Crude |  |  | Adjusted for ever taken a smear, years since training and current role |  |  |
|  | OR | 95% CI | p | OR | 95% CI | p |
| HPV knowledge | 1.48 | 1.23 – 1.78 | <0.01 | 1.27 | 1.03 – 1.56 | 0.03 |
| Triage and TOC knowledge | 1.68 | 1.41 – 2.00 | <0.01 | 1.29 | 1.05 – 1.59 | 0.02 |
| HPV vaccine knowledge | 2.03 | 1.55 – 2.65 | <0.01 | 2.03 | 1.41 – 2.93 | <0.01 |

Feeling adequately informed was independently predicted by HPV vaccine knowledge after adjustment for the other predictors, while the association with HPV knowledge and Triage and TOC knowledge disappeared after adjustment. All three knowledge domains independently predicted confidence in answering patient questions, even after adjustment for the other predictors.

### Improving traning

Suggestions for how training might be improved were provided by 36 respondents (15.7%). They wanted regular updates, more training sessions and several people felt that online training and other online resources such as research, frequently asked questions and updates would be useful. A request for specific advice that should be provided to parents and simple information sheets for both primary care and patients was suggested. There were also requests to widen the provision of training beyond practice nurses to all healthcare providers, specifically including GPs, independent vaccinators and Public Health Nurses delivering the School Based Immunisation programme.

## Discussion

Although mean knowledge levels for HPV and the HPV vaccine were reasonable (with each subset of questions yielding a mean percentage correct score of between 88% and 85%, respectively), only 25.2% and 44.3% of respondents scored 100% in each category, respectively. Knowledge about triage and test of cure was lower (mean percentage correct score of 74%) and only 9.1% of individuals correctly answered all the answers in this section. Some questions in each section revealed worrying knowledge gaps.

General HPV knowledge was the highest overall but around a quarter of respondents were unaware that having sex at an early age increases the risk of getting HPV. A quarter of respondents were also unaware that HPV is so common that most sexually active people will be exposed to it in their lifetime. Research has shown that considerable stigma can be attached to a positive HPV test^7, 8^ and that a lower level of education can be associated with an increase in the negative emotions and stigma that patients experience^8^. Therefore, it is vital that clinical staff are aware of the widespread nature of the virus so that they can reassure patients and reduce stigma. Furthermore, they need to ensure that when terminology such as ‘high risk HPV type’ is used, that this does not further increase patients’ stigma. A third of participants did not know or were unclear that HPV does not usually need treatment. This lack of knowledge has the potential to spread misinformation and cause confusion among patients as they seek treatment that is not available. Perhaps most worryingly, 14% of respondents either believed that HPV causes HIV/AIDS or were unclear that it did not.

In the triage and test of cure questions, while some questions were generally answered accurately, some questions revealed uncertainty and a lack of understanding of the current guidelines. For example, fewer than half of the respondents knew that not all cervical samples showing mild cellular changes are tested for high-risk HPV (only those for women aged 30 and older are tested for HPV under the current NZ guidelines). In addition, almost a quarter of respondents did not know or were unsure that a negative HPV test means that a woman is at low risk from cervical cancer. This uncertainty is likely to be problematic when primary HPV testing is rolled out. Unlike the cell changes that are screened for currently, primary HPV screening is about identifying a woman’s risk factors. Health professionals will need to be confident in talking with women about what their positive test result means. The test of cure questions were also correctly answered by fewer than three quarters of the respondents.

HPV vaccine knowledge was better, although there was confusion about the protection provided by the vaccine, with a fifth of respondents unclear that the HPV vaccine protects against most cervical cancers (HPV types 16 and 18 are responsible for 70% of all cervical cancers). A third of respondents were unsure or believed that the vaccine does not protect against genital warts. However, all HPV vaccines used in NZ offer protection against low-risk HPV types 6 and 11, which are responsible for 90% of genital wart infections.

Having ever taken a smear independently predicted Triage and TOC knowledge, as did the number of years since training. The number of years since training also predicted HPV knowledge. This pattern was repeated for HPV vaccine knowledge, however the results were not statistically significant, which could be related to the smaller scale on which HPV vaccine knowledge was measured, resulting in less variability in scores and making it difficult to detect small effects. There was some evidence of current role being associated with knowledge scores, with colposcopy workers being more likely to have higher Triage and TOC scores and laboratory staff and other workers being less likely to have higher HPV vaccine knowledge scores. However, some of these groups had very small numbers. The vast majority of respondents indicated that they would recommend the vaccine and they also favoured vaccinating boys and men, with only one individual indicating they would not recommend vaccinating boys and men. This is particularly reassuring since NZ made the vaccine available to boys from January 2017.

### How do NZ healthcare practitioners compare?

Research has been conducted in other countries with HPV vaccination programmes to explore healthcare practitioner knowledge about HPV and the vaccination. The studies reviewed below all reveal that, consistent with our NZ results, health care practitioner knowledge about HPV and the HPV vaccination is frequently incomplete.

An evaluation of knowledge about HPV and HPV vaccination for GP practice nurses in Leicestershire in the UK, where the vaccination has been administered through the NHS since 2008, found that although general HPV knowledge scores were quite high, there were specific gaps or weaknesses in knowledge^12^ as identified above. There were also gaps in their knowledge about current NHS processes around HPV triage and test of cure. For example, the role of HPV testing post-treatment (TOC) was misinterpreted, with only 66% acknowledging that all normal, borderline nuclear and mildly dyskaryotic samples are tested for high risk HPV post-treatment. Not all nurses felt adequately informed about HPV and a need to improve the provision of training was identified.

Nilsen et al explored knowledge of and attitudes to HPV infection and vaccination among public health nurses and GPS in Northern Norway in 2010, one year after the HPV vaccination was introduced for 12 year-old girls in Norway^14^. Knowledge of HPV infection, vaccine and cervical cancer was measured with 7 open-ended questions (e.g. what is the lifetime risk of a sexually active person getting HPV?). The percentage of GPs getting each question correct ranged from 26-55% while for the nurses it was 35-86%. Self-reported knowledge was considerably higher than actual knowledge. Only 47% of respondents knew that HPV infection is a necessary cause of cervical cancer.

In Malaysia there has been a school-based HPV vaccination programme since 2010. Jeyachelvi et al conducted a survey to explore HPV and HPV vaccination knowledge and attitudes in primary health clinic nurses who run the vaccination program in Kelantan, Malaysia^15^. Nurses were given 11 questions to assess their knowledge. The mean score was 5.37 with the minimum score being 0 and the maximum being 9. No question was answered correctly by more than 87.3% of respondents and the poorest question (External anogenital warts increase the risk of cancer at the same site where the warts are located. True/False) was answered correctly by only 10.6%.

Rutten et al conducted a survey exploring clinician knowledge, clinician barriers and perceived parental barriers to HPV vaccination in Rochester US^16^. They found that greater knowledge of HPV and the HPV vaccination (assessed together using an 11-item scale) was associated with higher rates of HPV vaccination initiation and completion of the 3-dose vaccination schedule, suggesting that knowledge is important in order to effectively promote HPV vaccination in addition to reducing stigmatising attitudes of clinicians as mentioned in the Introduction.

### Education needed

Our results suggest that education about HPV and particularly the use of HPV testing in the screening programme and test of cure process is urgently needed to address some worrying gaps in knowledge. This is especially important since further changes to the screening programme are due to be implemented, with draft primary HPV screening guidelines recently out for consultation^17^. As other countries also start to roll out primary HPV screening, the success of primary screening engagement in NZ and the rest of the world may well rest upon the level of knowledge of those people responsible for implementing it.

The need for education indicated by the knowledge scores was further reinforced by the fact that over a third of respondents did not agree that they felt adequately informed about HPV and that being adequately informed and feeling confident in responding to patients’ questions were both associated with knowledge. Suggestions for training were proposed by some of the respondents. One promising suggestion, which was also proposed by UK practice nurses,^12^ was for online training. This would provide a low-cost way to update changes to the vaccination and/or screening programmes and guidelines in a format that would be easily accessible to many staff whilst requiring relatively little time commitment to complete. HPV vaccination online training was developed by the Immunisation Advisory Committee (IMAC) and was released in August 2017^18^. This may address some of the knowledge issues associated with vaccination identified in this study, but additional online training regarding screening and test of cure is needed.

Our survey is the first to be conducted in NZ that explores health practitioner knowledge and understanding about HPV, the vaccine and the role of HPV testing in the cervical screening programme and it contributes to the international picture about HPV knowledge that is emerging. It is evident from our findings and those from other countries, that more education is required to ensure that misinformation, stigma and widening inequalities do not inadvertently result from interactions between health practitioners and the public.

## Data Availability Statement

The information sheet, survey and data are deposited publicly on the Open Science Framework website and can be accessed here: osf.io/ub7g2, DOI 10.17605/OSF.IO/UB7G2.

## Acknowledgements

We would like to thank the following individuals for assistance with data collection:

Jane Grant – Cervical Screening Nurse Specialist, Metro Auckland Cervical Screening Coordination Service, Auckland and Waitemata DHBs
Lucina Kaukau - Cervical Screening Nurse Specialist, HPV Self-Sampling Feasibility study for Maori women, Research Nurse.
Lisbeth Alley – Programme Manager, Immunisation. Auckland and Waitemata DHBs
Pam Hewlett – Women’s Health Manager. Auckland and Waitemata DHBs

## Author Contributions

Conceived and designed the study: SS, CB. Collected the data: KB, CB, HD. Analyzed the data: SS, HD. Wrote the paper: SS, KB, HD, HP, EM, JD, CB.

## Ethical Approval

Ethics approval was granted by the Massey University Ethics Committee 4000015595

The project was registered with Waitemata DHB localities (Reference number RM13518). Both Waitemata and Auckland DHB confirmed that locality authorisation was not required as the research was carried out in community health care settings.

